# Covariations of cerebral blood volume and single neurons discharge during resting state and behavioral visual cognitive tasks in non-human primates

**DOI:** 10.1101/2022.06.20.496840

**Authors:** Julien Claron, Matthieu Provansal, Quentin Salardaine, Pierre Tissier, Alexandre Dizeux, Thomas Deffieux, Serge Picaud, Mickael Tanter, Fabrice Arcizet, Pierre Pouget

## Abstract

To better understand how the brain allows primates to perform various set of tasks, the ability to simultaneously record the activity of the brain at multiple temporal and spatial scales is challenging but necessary. In non-human primates, combined fMRI and electrophysiological recordings have not disentangle the contributions of spiking activity to the neurovascular response. Here, we combined functional ultrasound imaging (fUS) of cerebral blood volume (CBV) and recording of single-unit activities (SUA) in visual and fronto-medial cortices of behaving macaques. We computed task-induced and SUA-induced CBV activation maps. We demonstrate that SUA provides a significant estimate of the neurovascular response below the typical fMRI voxel spatial resolution of 2mm^3^. Furthermore, our results also show that single unit and CBV activities are statistically uncorrelated during the resting states but correlate during behaving tasks. Conversely, during the resting states, CBV activities across known connected brain areas are correlated but decorrelate at the onset of the tasks as expected if participating in the default mode network (DMN). These results have important implications for interpreting functional imaging findings collected with fMRI or fUS while one constructs inferences of spiking activities during resting-state or while primates perform tasks.

---

Awe-inspiring progress was enabled by novel technologies for imaging the entire brain at microscopic scale and a very high temporal resolution to study neural functions in flies, fishes or even rodents^1–3^. In large animals, the pioneering studies from Logothetis and colleagues showed that the impulse response of the neurovascular system correlates better with LFPs compared to multi-unit activity, suggesting that the BOLD signal predominantly reflects the input and intra-areal processing^4^. Similarly, other studies found that BOLD fluctuations loosely relate to action potential frequencies but are rather coupled to LFP oscillations in cats^5,6^ and in non-human primates^7,8^. However, other studies have shown that the BOLD signal provided a reliable measure of firing rates in cats^9^, non-human primates^10,11^ and humans^12^. For example, a recent study showed that BOLD-derived population receptive fields (pRF) were more like pRFs based on multi-unit activity compared to LFPs^13^. Others demonstrated that visual stimuli-induced early variations of oxygen are reliable estimates of spiking activity. Moreover, hemodynamic activity is also dependent on other non-neural fluctuations^14^. Finally, optogenetic driving of neuronal activity generates BOLD signals that correlate with firing rate in rodents^15,16^. This disparity of results can be resolved using an imaging technique with higher spatial resolution than fMRI to investigate the contribution of single unit spiking to the neurovascular system.

Functional ultrasound imaging (fUS) is an innovative imaging technique which can provide whole-brain maps of neurovascular activity changes^17^ even in deeper regions (up to 1.5 cm) with an approximately 5 to 10-fold better spatiotemporal resolution and sensitivity than fMRI (100 µm, 1Hz, even if uniplanar fMRI sequences allow the same temporal resolution, the spatial resolution is unachieved with typical fMRI sequences). Like fMRI, fUS technique relies on the neurovascular coupling of brain activity but unlike BOLD signals, fUS measures changes in the Cerebral Blood Volume (CBV) within microvessels using ultrafast doppler^18^ combined with spatiotemporal clutter filtering^19^. fUS technique requires no large magnetic machinery and so has been applied in freely moving small animals using miniature, head-mountable transducers^20^. Several studies used fUS to investigate sensory processing in rodents^21–25^ and in behaving primates^26–28^. Besides, an increasing number of studies have demonstrated that fUS can monitor brain activity in humans during surgeries^29^, but also through the skull bone in neonates^25,30,31^, or even adult patients^32^. Further knowledge of the contribution of neuronal activity to the CBV variations in humans and close species is therefore important for future interpretations of such basic and preclinical studies. To our knowledge, fUS and recording of nearby spiking activity has been achieved in rodents only^33,34^. However, the rodent model does not allow complex behavioral studies and brain architecture is quite far from that of humans, and therefore, NHP model might allow to transpose the ins and outs more easily to the human brain. One recent publication concomitantly performed neural activity recordings and fUS for the first time in mice^35^. Their results show that CBV variations match the smoothed firing rates of nearby neurons, especially inhibitory, in the visual cortex and the hippocampus. A direct comparison between fUS imaging signals and spiking activity has not been achieved in behaving primates. Furthermore, such comparison permits to characterize a bit more the nature of resting state in primates, which has been extensively studied in humans^36,37^ but rarely to a coupled neuronal/vascular point of view. Consequently, new protocols are needed to achieve simultaneous recordings of CBV and neural spiking in behaving primates. Most fUS studies have assessed the significance of CBV variations^17,21,27^ sometimes in comparison with spiking activity^33^, using covariate statistical analysis. To go a step further, we implemented from the fUS signal a Generalized Linear Model methods (GLM) estimate allowing us to separate stimulus-induced, neural spiking and noise signals. This approach facilitates direct comparison of our results with those from fMRI studies.

Using a new protocol including ultrafast and high-resolution ultrasound imaging techniques (fUS) combined with commonly used electrophysiological systems in awake non-human primates. We recorded both fUS and single unit activities in visual cortex and SEF for Monkey L and Monkey S respectively while animals performed passive fixation and saccade tasks. We chose to focus on two different cortical areas to generalize the relationship between neural spiking and CBV variations. Our method permits online micro-electrodes position tracking and the recording of electrophysiological and CBV signals in awake behaving animals. We computed stimuli-induced and spiking-induced activation maps using a GLM. Our results demonstrate that individual unit activities provide a significant estimate of the neurovascular system response.

Our first main result is that we could use our imaging method to perform precise online tracking and guidance of micro-electrodes. The chambers were located above the Supplementary Eye Field (SEF) and the visual cortex V1 for Monkeys S and L, respectively (Fig. 1.a and Fig. 1.b). The adaptor was designed and 3D printed to allow the technical challenge of co-recording nearby spiking signal and CBV variations of the imaging plane. By slowly inserting the microelectrode into the brain, we detected on the image a precisely localized high rise in the Power Doppler in the region of interest at z=5mm (Fig. 1c), indicating that the tip of the electrode reached the imaging plane. When we inserted the electrode further, the high-Power Doppler signal extended vertically downward, allowing us to detect the electrode progression through the thickness of the imaging plane from z=5mm to z=12mm (Fig 1.c). Indeed, we noticed a higher power doppler signal localized at the position of the tip of the electrode (Fig 1.c) characterizing its motion during the insertion. Thus, our method permits precise online tracking of micro-electrodes in the brain in awake behaving primates.

**Figure 1:**
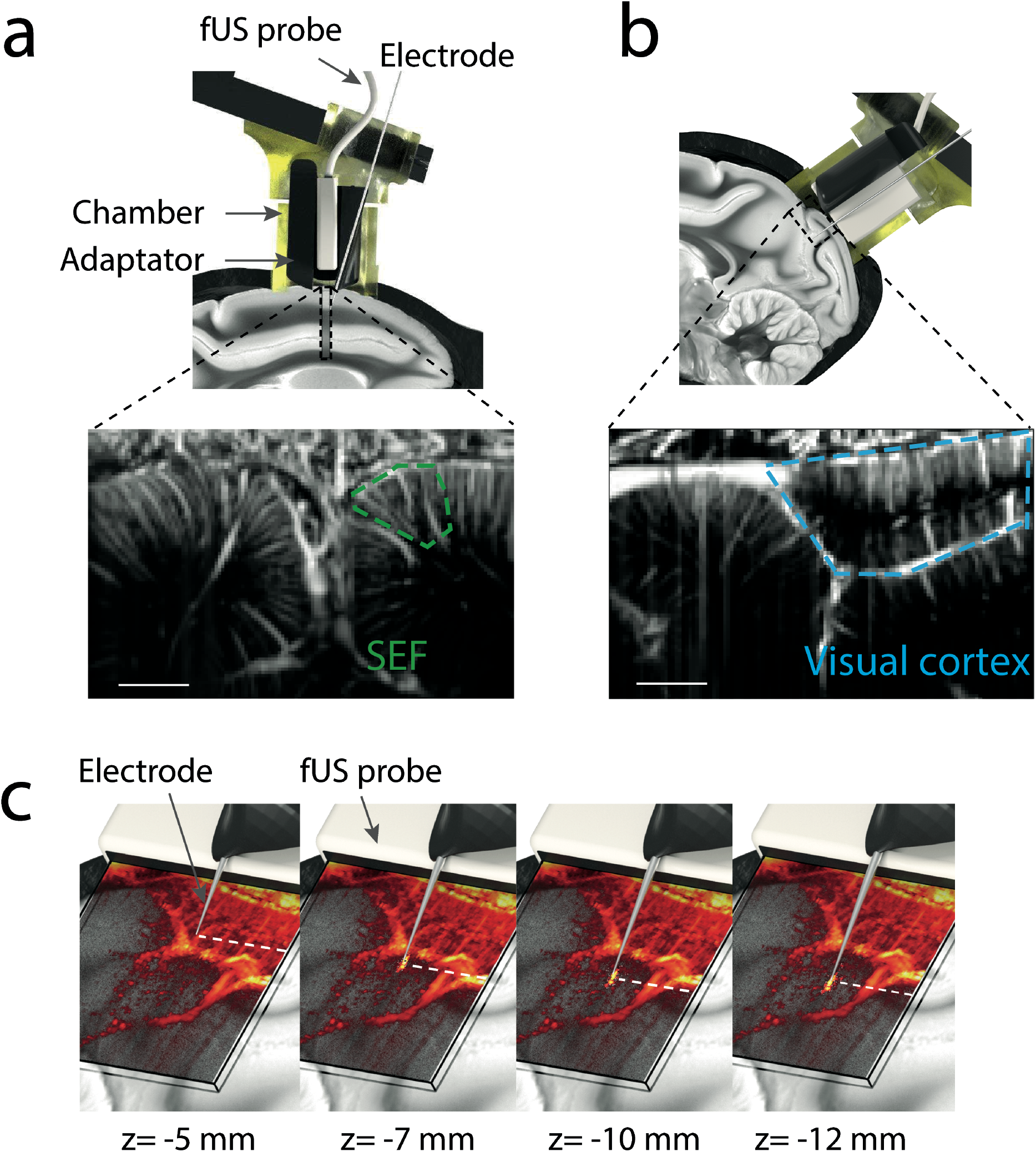
experimental setup. **(a)** Co-localization adaptor. We used this set-up on both animals; here it is illustrated for the recording of SEF activities. Top panels: a custom cylindric adapter allowed the placement of a custom fUS probe (128 elements, 15 MHz, 100 × 100 μm² of spatial resolution) into the recording chamber. A micromanipulator was screwed on top of the adaptor and controlled the insertion of a single-channel electrode through the cortex via a tilted guide tube so that the tip of the electrode was placed in the fUS imaging plane. Bottom: fUS of the SEF. **(b)** Co-recording adapter. We used this set-up on both animals; here it is illustrated for the recording of the visual cortex activities. Same as the co-localization adapter except the single-channel electrode is inserted parallel to the fUS imaging plane. Spatial scale bar: 2mm **(c)** Online tracking of micro-electrodes. fUS of the region of interest (visual cortex here) was performed while inserting a single-channel electrode through the brain tissue. A high Power Doppler signal was detected when the tip of the electrode reached the fUS imaging plane at z=5mm. This high Power Doppler signal extended downward when further inserting the microelectrode (z=7mm, z=10mm and z=12mm, respectively).

Having established a method to place micro-electrodes accurately in or near the fUS plane, we concomitantly recorded CBV and single unit activity while the animals performed behavioral tasks. Monkey S performed a visual stimuli-guided pro-saccade or anti-saccade task with equal proportion for both conditions (Fig 2.a., see Methods). Monkey L performed a passive fixation task during which a peripheral visual stimulus was presented (Fig. 2.b.). We used a repertoire of different visual stimuli (see Methods) to generate different activation maps of the visual cortex. For most sessions, a single stimulus was presented. For others, hemi-concentric bands of varying eccentricities or squared-shape gratings with varying spatial frequencies were used (Fig. Supp. 2). For Monkey S, we recorded a total of *n*=18 fUS imaging sessions and for Monkey L, *n*=79 fUS imaging sessions (53 in which we also co-recorded single-units). For V1, we assessed that there were no significant differences of maxima, minima and their respective timings between the mean CBV responses of each stimulus type (Kruskal-Wallis, p>0.05, Fig S1). Therefore, we generated a single hemodynamic response function (HRF) by averaging the mean CBV response over sessions and computing the gamma inverse function that fit best. Hence, we computed the activation map induced by all stimuli presentations (independently from their nature) using the GLM approach. We did not focus on the spatial regionalization of the activation among stimuli, but see the works of Blaize and colleagues for detailed retinotopic maps in the visual cortex^27^. For further analyses about the dynamic of activation between pro- and anti-saccade, see previous works of Claron and colleagues^38^. In our model, we took as input the spatiotemporal Doppler images and took a stimuli-induced regressor (Fig 2.c. and Fig 2.d). Briefly, for each pixel, all visual stimulus presentation timings were convoluted with the fUS-determined HRF of the investigated cortical area to obtain a CBV signal prediction. From the fit between these signal predictions and the real CBV variations Z-score and p-value map were obtained (Fig. 2.e for SEF and Fig 2.g for V1).

**Figure 2:**
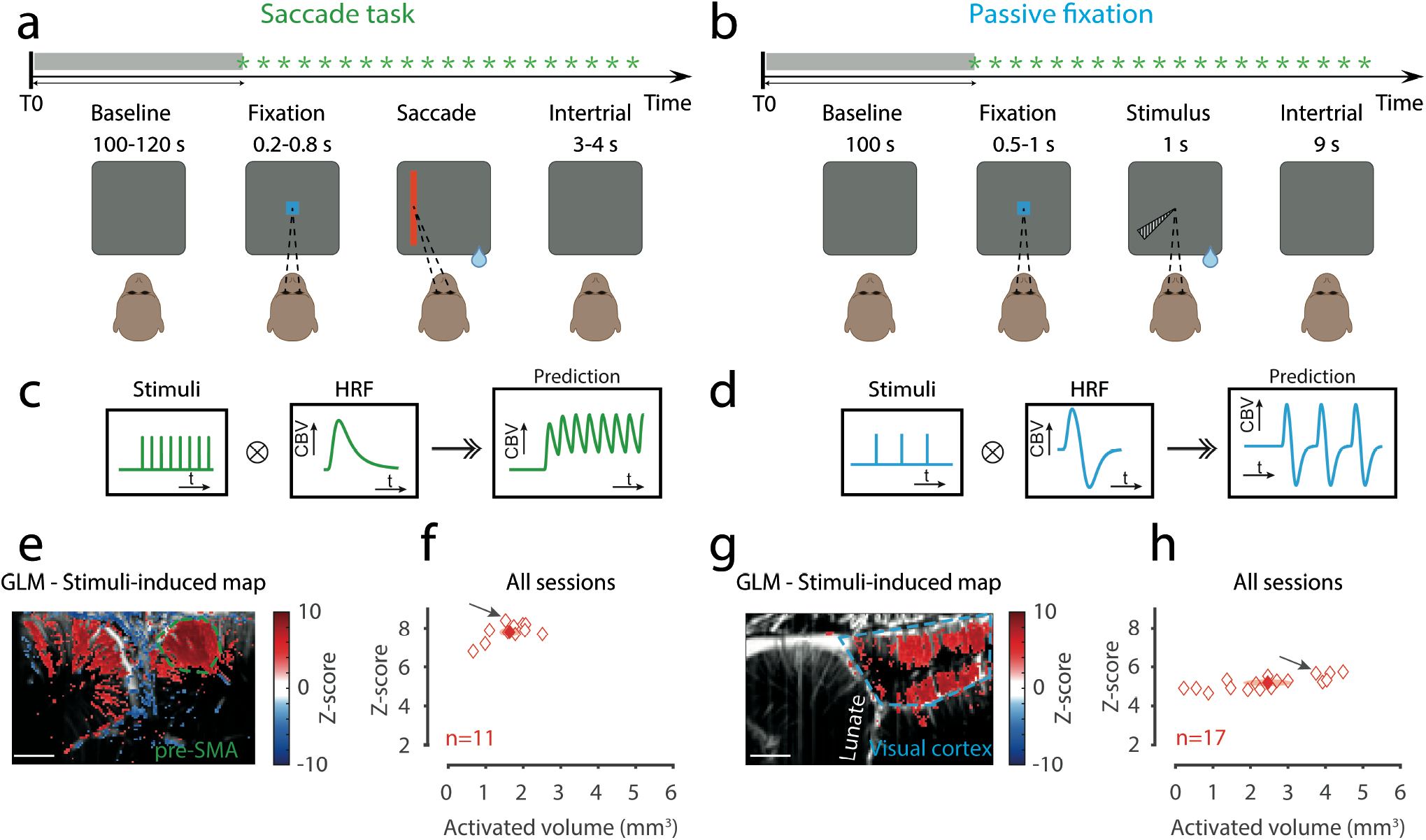
Task-related CBV responses. **(a-b)** Behavioral tasks. Both tasks began with a baseline period (no stimulus). (**a)** Saccade task. Following an initial fixation of a central spot, the animal had to perform a saccade or anti-saccade depending on the cue that appeared on screen (vertical or horizontal bar, respectively). (**b)** Passive fixation task. Following an initial fixation of a central spot, the animal had to maintain its gaze during 1s while a visual stimulus (see Methods for the different stimuli) appeared on screen. (**c-d)** Task-based Generalized Linear Model (GLM). For each animal and cortical area (blue: SEF, green: visual cortex), we computed a behavior-based ΔCBV prediction by convoluting the stimulation pattern (timings of cue presentation) with the hemodynamic response function (HRF). We then obtained activation maps that represent the Z-score for all voxels above a significance threshold (p<0.001 after Benjamini-Hochberg correction). (**e)** and (**g)** respectively display representative activation maps of the SEF and the visual cortex. Coloured dashed lines represent the regions of interest. (**f)** and (**h)** display scatter plots of the median Z-score of activation of each session against its volume for SEF and visual cortex, respectively. Plain symbols display mean data. Ellipses represent the SD in each dimension. Spatial scale bar: 2 mm.

The activation maps show the Z-score of all pixels in the images with a p-value < 0.001 (after Benjamini-Hochberg correction to decrease false discovery rate). Fig. 2.e shows a representative activation map induced by the presentation of both pro-saccade and anti-saccade stimuli for correct trials for Monkey S (see Methods for GLM application). We detected significant pixels in both hemispheres although the right SEF (dashed green line) displayed more significant pixels, corresponding to an activated volume around 2 mm^3^. All sessions were balanced in terms of ratio pro-v.s. antisaccades and left vs right. Fig. 2f represents the population analysis of median z-scores and volume of activations in SEF for sessions in which activation was detected in the region of interest (*n*=11). The mean stimuli-induced volume of activation in the SEF was 1.64±0.54 mm^3^. Figure 2.g shows a representative activation map in the visual cortex induced by the stimulus presentation for correct trials for Monkey L. We detected significant pixels in the superficial and deeper layers of the region of interest (V1 and V2, respectively) corresponding to an activated volume of 3.7 mm^3^. Fig. 2h represents the population analysis of median z-scores and volume of activations in the visual cortex for sessions in which activation was detected in the region of interest (*n*=17). The mean stimuliinduced volume of activation was 2.5 ± 0.7 mm^3^.

For Monkey S, among *n*=18 imaging sessions, we recorded *n*=18 single-units. Among this pool of 18 neurons, we obtained *n*=11 single-units displaying significant response locked to the saccade execution. For Monkey L, among *n*=53 imaging sessions, we recorded *n*=17 single-units displaying significant response locked to the stimulus presentation. We then investigated the neurovascular response associated with isolated single-unit activities. To visualize the dynamics of the CBV responses, we plotted the averaged variation of CBV over time for all correct trials and for all pixels of each region of interest (delimitated by dotted lines in Fig. 2). The SEF CBV variation rose after a dip synchronized with the stimulus presentation (t=0s) and reached a plateau around 3% 1.8s later, before decreasing back to its initial value after 4s (Fig 3.a., bottom panel, red curve). Here, during the same example session, we recorded the activity from a single unit located in the right SEF. The raster plot (Fig. 3.a, top panel) and the spike density function (Fig 3.a., bottom panel, blue curve) show a transient increase of the firing rate (up to 30 Hz) that corresponds to the saccade execution, followed by a less active period. Note the different kinetics between the fast neuronal activation and the slower neurovascular response; the peak of the CBV response occurred 1.6 s after the firing rate peak.

**Figure 3:**
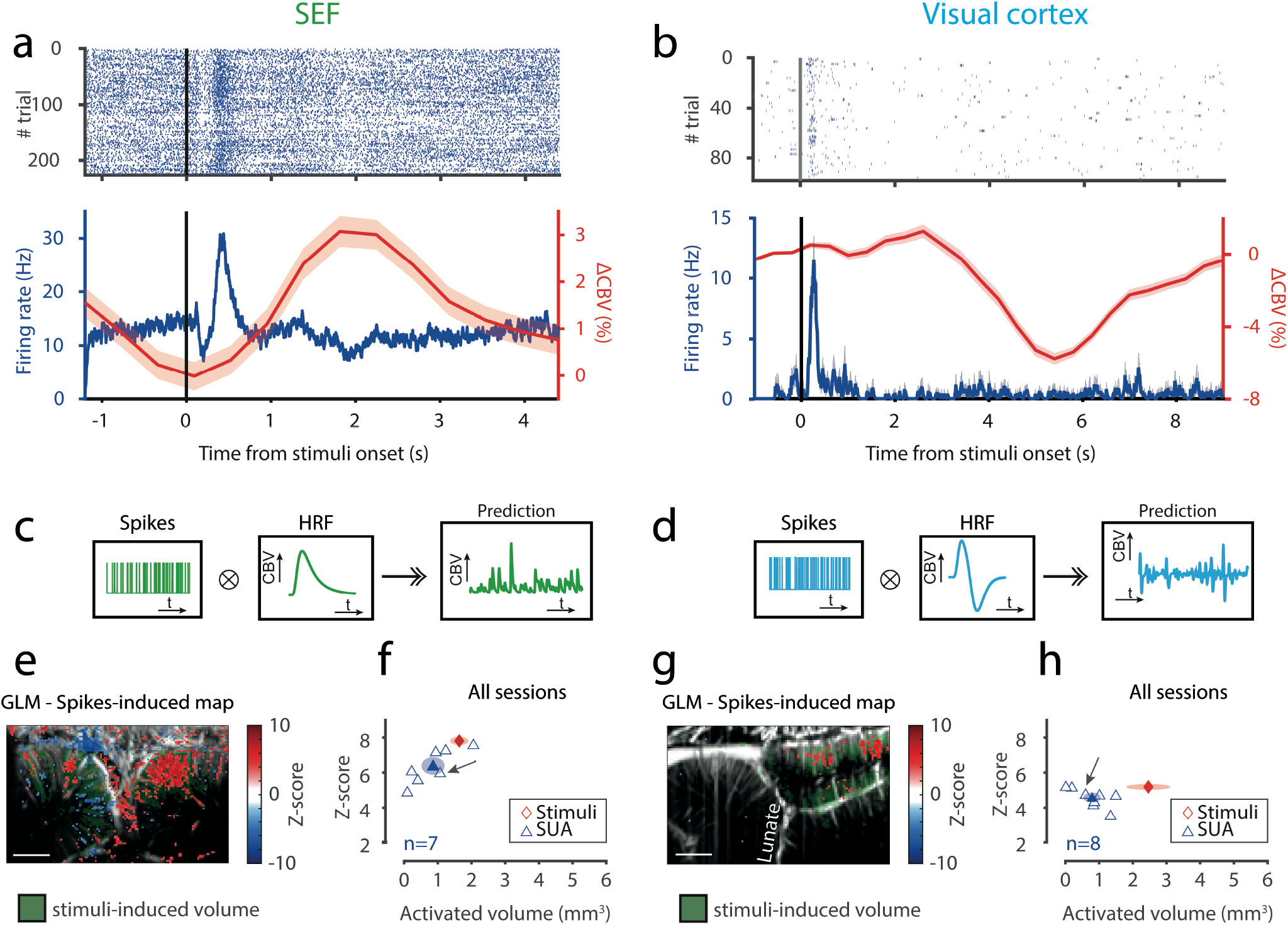
Spike’s contribution to the CBV response. **(a)** Top: raster plot of the SEF single-unit that was co-recorded with fUS, for correct trials only. Bottom: spike density function (SDF, blue curve) of the same single-unit as above and ΔCBV from the ROI delimited in the SEF map in (e). **(b)** Same but for the visual cortex activities extracted from the same example session than the visual cortex activation map in (g). t=0 refers to the cue presentation. Data presented are mean +/-s.e.m. **(c-d)** GLM based on spike activities. Diagrams in each panel: same as Fig. 2. (c-d) but the stimulus pattern is replaced by the spikes onset. **(e)** and **(g)** Representative spike-induced activation maps of the SEF and the visual cortex, respectively. Coloured dashed lines represent the regions of interest. Stars represent the electrode insertion axis (distance *circa* 1mm tangent to the imaging plane). Green-colored pixels represent the behavior-induced CBV response (shown in Fig 2.(e) and (g)) which is more spread. (**f)** and (**h)** display scatter plots of the median Z-score of activation of each session against its volume for SEF and visual cortex, respectively. Task-induced and SUA-induced activations are represented by blue diamonds and red triangles, respectively. Plain symbols display mean data. Ellipses represent the SD in each dimension. Spatial scale bar: 2mm.

We next wanted to test whether the spiking activity recorded with the single electrode at a close location could contribute to the neurovascular response. To do so, we used another GLM approach based on the spike’s activity. Here, we convoluted for each animal the single unit spike onsets with the HRF extracted from the same region of interest (Fig. 3.c for SEF, Fig 3.d for V1) to obtain a CBV signal prediction. As previously, the representative co-activation maps show the Z-score of all pixels in the images with a p-value < 0.001 (before Benjamini-Hochberg procedure to decrease false discovery rate) obtained from the fit between the predicted signal and the real CBV variations (Fig 3.e). We also represented the task-related activities displayed in Fig 2.e for comparison (green transparent colored pixels). For Monkey S, the co-activation map showed a spikes-correlated neurovascular response in the SEF corresponding to a volume of ∼ 1.1 mm^3^ (Fig 3.g, left panel) compared to 2 mm^3^ with the GLM based on stimuli. Interestingly, the spread of this co-activation was more localized than the activation detected with the GLM approach based on the stimulation pattern. Furthermore, the spread of this correlated activation was constrained around the electrode recording site (indicated by the dashed white line). We repeated these analyses for the visual cortex that revealed a different pattern of CBV variation. Indeed, the CBV variation displayed a biphasic increase up to 1.3% (Fig3.b, bottom panel, red curve). Interestingly and unlike for the SEF response, we detected a large undershoot in the visual cortex neurovascular response (from 4 to 8s) before the signal went back to its initial value. The raster plot (Fig.3.b, top panel) and the spike density function (Fig 3.b, bottom panel, blue curve) of simultaneous single unit recording show a transient increase of the firing rate (up to 11.5Hz). Here, the neuronal activation was also faster than the neurovascular response with a difference of 2.45 s. We reported consistent results for Monkey L (Fig 3.g, right panel). As for Monkey S, the spread of the spikes-correlated activity was smaller (∼0.6 mm^3^, Fig 3.h) than the activation induced by stimuli onsets but was not particularly restricted around the electrode recording site. To conclude, these results demonstrate our method allows examining specific links between single-unit activities and cerebral blood volume variations in behaving non-human primates.

To assess that the single unit-induced activation is more localized than the stimuli - induced activation, we computed the volumes and median Z-scores of activations for each condition over our entire pool of data (Fig 2.f and Fig 3.f for SEF, Fig 2.h and Fig 3.h for V1). For Monkey S, we showed that the mean stimuli-induced volume of activation in the SEF is 1.64±0.54 mm^3^ whereas the SUA-induced volume of activation is 0.86±0.68 mm^3^. (Student T-test between the stimuli-induced- and the SUA-induced activities gives the former significantly higher than the latter, with p = 0.016, T-test is chosen as data are gaussian (Shapiro-Wilk test, p = 0.82 and p = 0.58 respectively) and F-test gives similar variances (p = 0.49)). Peak Z-score of GLM-calculated maps was found slightly higher for the stimuli-induced map (7.81±0.47) than the Z-score for the spikes map (6.39±1.00; p = 0.0084, Shapiro-Wilk test assures both Z-scores for stimuli-induced and SUA-induced are gaussian, F-test assures that variances are unequal). We confirmed these results in the second animal: for Monkey L, the mean stimuli-induced volume of activation of the visual cortex was significantly higher than the SUA-induced volume of activation (stimuli-induced: 2.5 ± 0.7 mm^3^, *n*=17 acquisitions; SUA: 0.8 ± 0.5 mm^3^, *n*=8 responsive units, p=1.3915.10^-4^, two-sample T-test with unequal variances as samples are gaussian according to Shapiro-Wilk test (p = 0.6064 and p = 0.8096 for stimuli-induced and SUA-induced, respectively) as illustrated in Fig 3.g for the example session. Interestingly, the median Z-score of stimuli-induced activation was also significantly higher than the median Z-score of SUA-induced activation (stimuli-induced: 5.19 ± 0.17, *n*=17 acquisitions; SUA: 4.58 ± 0.55 mm^3^, *n*=8 responsive units, p=0.0024, two-sample T-test as samples are gaussian according to Shapiro-Wilk test (p = 0.2882 and p = 0.4771 for stimuli-induced and SUA-induced, respectively) and have equal variances according to F-test (p = 0.0947). Altogether, these results demonstrate that single-unit activities partially contribute to the cerebral blood flow variations in behaving non-human primates.

With GLM analyses, we spatially characterized the relationship between neural spiking and CBV in two different cortical areas during behavioral tasks. One could also ask how both CBV and neural spiking are temporally linked. We next investigated how neural spiking contribution to the neurovascular activity changes between resting state (i.e. BL=baseline, 100 to 120s-long period before task onset) and the start of the task. We first represented display examples of CBV variations and SDF over one unique session for SEF and V1, respectively (Fig. and Fig. 4.b). We computed the moving non-overlapping cross-correlation across resting state and task between the fUS signal in the ROI and the SDF of the recorded single-unit. Fig. 4.c and Fig. 4.d. left graphs represent the mean and s.e.m. of these moving cross-correlations across sessions for SEF (n=9) and V1 (n=15, two sessions missing initial baseline), respectively. We quantified the average values of cross-correlation during resting state and task for SEF and V1, and for two control areas: median cingulate cortex (MCC) for monkey S and a 1×1 mm² squared ROI located at the bottom left corner of fUS images for monkey L (see the locations of these areas in the maps Fig 4.c. and Fig. 4.d). Interestingly, both in the SEF and in V1, there is no correlation between SDF and fUS signal during resting state (t-test comparison to null distribution, SEF : p = 0.96, V1 : p = 0.56) but these signals became correlated in SEF and anticorrelated in V1, respectively, after the task began (SEF r = 0.2 +/-0.03 s.e.m, p = 2.6e-3, V1 r = -0.06 +/-0.0096 s.e.m, p = 0.0204, t-test). The anticorrelation in V1 might be due to the undershoot of the HRF in this cortical area. Finally, we computed the correlation between CBV variations of different areas of the brain, both during resting state and during task in order to characterize the vascular component of the default mode network and the variation due to the beginning of a behavioral task. In the median wall of primate brain, we observe a correlation between SEF and the MCC during resting state (Fig 4.c, right panel, r = 0.42 +/-0.07 s.e.m., p = 3.4e-4 t-test comparison to null distribution) and a decrease of this correlation between the SEF and the MCC during the task (r = 0.15 +/-0.11 s.e.m., p = 0.20 compared to null distribution, p = 0.03 compared to resting state, t-test) suggesting a vascular component of the default mode network and an alteration of vascular co-oscillations during active behavior whereas neuron spiking and vascular activities synchronize during behavioral tasks. Similarly, in the visual cortex, the correlation between V1 and V2 during the baseline is higher during the resting state (Fig. 4.d, right panel, r = 0.66 +/-0.019, p = 5.0e-11 compared to null distribution) than it is during task (r = 0.39 +/-0.083, p = 3.66e-4 compared to null distribution, p = 0.0038 compared to resting state, t-test).

**Figure 4:**
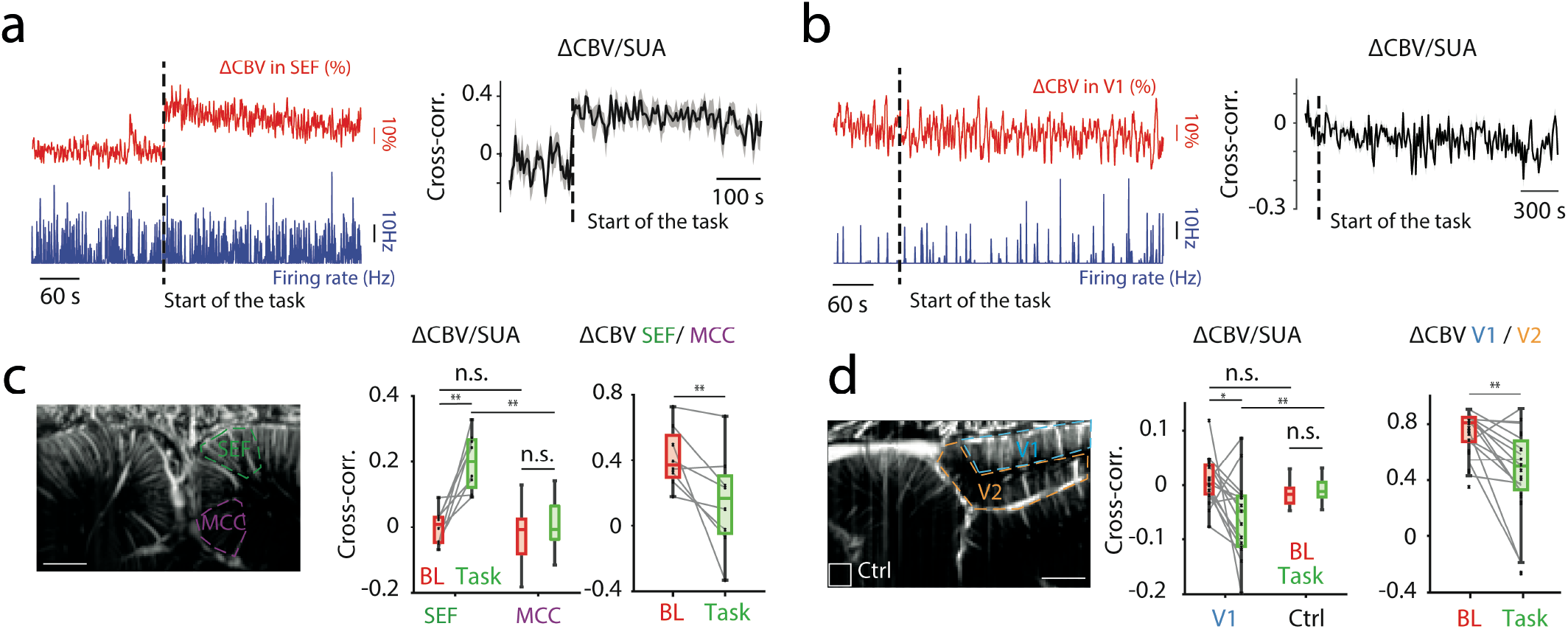
Intra- and inter-areas cross-correlation of neural spiking and CBV variations shift when behavioral task is engaged. **(a)** Left panel: example of CBV variations (red curve) and SDF (blue curve) during session for SEF. Right panel: mean sliding cross-correlation between CBV variations and SDF in SEF across all sessions (*n*=9). Grey shaded area represents s.e.m. **(b)** Same as (a) for V1 (*n*=15 sessions). **(c)** Left graph: mean cross-correlation between CBV variations and SDF during resting state (i.e., initial baseline (BL) of the recording session) and task for SEF and median cingulate cortex (MCC). Right graph: cross-correlation between CBV variations of SEF and MCC during resting state and task. Map: locations of the areas used before. **(d)** Same as **(c)** for V1 and V2. Control area is a 1×1 mm² squared ROI located at the bottom left corner of fUS images for monkey L.

We have developed a methodology and associated setup to record simultaneous CBV and electrophysiological signals in awake behaving macaques. Our method offers first to report online tracking of the microelectrode localization in the brain of behaving non-human primates. The spatial resolution of ultrasound imaging offers an unprecedented ability to co-register micro-electrode and micro-sensor and potentially any rigid pipettes around 100 µm wide within the Ultrasound imaging referential frame. Further studies might combine our method to localize viral injections’ pipette and may help quantify the spread of compound perfusion during surgeries with a deeper field of view than allowed by optical imaging^39^, possibly allowing injections in subcortical areas. Functionally, fUS and electrophysiology do not seem to interfere with one another, or at least we were not able to measure the interference between them both, thus the coupling of the techniques is far easier than fMRI, optical imaging, and electrophysiology in awake behaving primates. Previous studies have found that fUS could be combined with acute or chronic EEG recordings in awake rodents^40,41^ and multi- or single-unit recordings in anesthetized or awake rodents^33,34^. These pioneering studies in awake rodents have shown a marked change of fUS activity at the initiation of tasks but were limited to explore sensory or cognitive functions in deep cortical folded brain areas. Our study demonstrate that our methodology allows using standard protocols to record simultaneous CBV and electrophysiological signals in awake behaving macaques without movements or electromagnetic artifacts. For the statistical assessment of CBV, some of these fUS studies used correlation methods of time evoked variations of CBVs compared to the more standard protocols used in other imaging techniques such as fMRI. In this study we report fUS analysis based on GLM standard statistical assessment in NHP^42^. It depicts for each voxel associated time-series Y, as the linear combination of regressor of interest (e.g., oculomotor or visual tasks) including spiking activity of our awake behaving primates. This analysis proved the high sensitivity of our recording method and showed that brain activity could be investigated with minimal adaptation for different brain areas or electrophysiological systems (SEF and V1) in awake behaving primates. We hypothesized that single unit-induced activation is more localized than the stimuli-induced activation based on the significantly activated area estimated from the thresholded Z-score maps. This assumes that there is no large difference in the statistics between both maps so that thresholding does not artificially reduce the activated area size. This seems to be generally the case as both maps are constructed from the same acquisitions and parameters.

The relationship between both vascular and neuronal signals is still debated. Recent studies report striking inconsistencies between the fMRI and electrophysiological signals^43^. Previous fMRI studies have shown that the neurovascular activity (BOLD signal) provided a reliable measure of multi-unit activities^9–11^. Conversely, other studies have reported a poor correlation between multi-unit activities and the BOLD signal^5,7,8^. Importantly, we demonstrated here the cortical volume in which SUA correlate significantly with CBV variations is smaller than the cortical volume in which task events correlate with cue presentation timings. Indeed, few works have spatially characterized spikes’ contribution to the neurovascular system. It has been shown that blood flow variations in the olfactory bulb of rodents show a similar linear relationship with locally measured neuronal Ca^2+ 22^. A recent publication showed that neural signals from the visual cortex and hippocampus in mice, and more particularly firing rates from inhibitory neurons, also correlate with fUS imaging signals in mice. Co-activation maps from an fMRI study carried out in human subjects^12^ have shown that SUA correlation with BOLD signal in Heschl’s gyrus is more localized than the activity predicted by LFP, although this result was more qualitative than quantitative. However, the authors tested the correlations between SUA recorded in a first group of patients and the BOLD signals from other patients, whereas we perform here an intra-individual analysis. At least four hypotheses have been discussed that can explain the discrepancies between the two LFP and SUA methods: (1) the BOLD signal follows local field potential (LFP) signals closer than spikes^4^, (2) the BOLD signal is reflecting electrophysiological signals that are occurring later due to feedback delay^44^, (3) the BOLD signal is more sensitive than traditional electrophysiological methods due to massive pooling by the hemodynamic coupling process^45^, and finally (4) there is no real inconsistencies, and instead, small but reliable effects on firing rates may be obscured by differences in experimental design and interpretation of results across methods and statistical analysis^46^. Our main result demonstrates that a fifth explanation prevails. Indeed, fUS imaging enables us to show that the cortical volume in which SUA correlates significantly with CBV variations is smaller than the cortical volume in which task events correlate with cue presentation timings. We demonstrated this in two behaving non-human primates’ primary visual and fronto-medial cortices. It consequently indicates that single-unit activities correlate with CBV variations recorded if and only if the blood dependent signal is extracted at sufficiently high spatio-temporal resolutions. As this is the case in fUS imaging, we showed that single unit activities provide a significant estimate of the neurovascular response with an activated area smaller than usual fMRI spatial resolution. Thereby, fUS seems a technique of choice in order to study neurovascular coupling in the brain of awake behaving primates because of a higher spatial resolution (∼200 microns).

fUS imaging in awake behaving NHP allowed us also to study the difference of neurovascular responses between resting state and task. Resting state is believed to be the reflection of brain activity during rest^47^ but it is still unclear if this vascular activity, measured with fMRI, is the reflection of the neuronal activity, even though some studies use fMRI combined with EEG or MEG to assess link between them both^36,48–50^. We demonstrated in this study that CBV and single unit activities are uncorrelated during resting state and become correlated during various tasks. Furthermore, we found that CBV activities across known connected brain areas are correlated but decorrelate at the onset of the tasks as expected if participating in the default mode network (DMN). Therefore, BOLD measured in fMRI, as well as CBV measured with fUS, during resting state might not much reflect the neuronal activity but basal metabolic changes through glia and astrocytes or intrinsic connectivity through slow synchronized neuronal oscillations across distant brain areas.

With the current 2D visualization, displacement of the probe is required to map a large volumetric area. The 2D view remains one intrinsic limitation of our method to investigate the 3D brain activity. However, ongoing development to perform direct 3D ultrasound imagery with micro-probes would certainly offer new options soon as it is already done in rodents using 3D array probe^51^ or row-column addressed probes^52^. Finally, the use of multi-contact electrodes possibly introduced over an entire cortical region will allow the study of the dynamics and propagation of the relationship between electrophysiological signals and the vascular response. Within various parts of cortical areas of primates, the use of such a technique could undoubtedly allow the study of electrophysiological and vascular responses in layers and thus succeed in studying neuro-vascular coupling in primates at a spatial and temporal resolution never achieved.

## Data availability

All data underlying the figures are freely available on the following link: https://osf.io/v9xgb/

## Author contributions

J.C. and P.T. designed the coupling system. J.C, M.P and Q.S trained monkeys and acquired data. J.C. and M.P. analyzed data. A.D. helped for the figures. T.D., S.P., M.T., F.A. and P.P. supervised this project. Abstract, core text as well as figures were done by J.C., M.P., F.A and P.P.

## METHODS

### Animal model and behavioral data

All experiments were ethically approved by the French “Ministère de l’Education, de l’Enseignement Supérieur et de la Recherche” under the project references APAFIS #6355-2016080911065046 and #9013-2017021515254591. Functional data were acquired from two captive-born rhesus monkeys (Macaca mulatta), S and L, each one trained to perform different visual tasks.

Monkeys were seated in a primate chair (Crist Instruments) with their head fixed and placed in front of a cathodic computer screen, 58cm away in a darkened booth. Mean screen luminance was controlled (1.15 mW.cm^-2^). Eye position of the primate was monitored at 1 kHz using an infrared video eye tracker (Eyelink 1k, SR-Research), which enabled live control of the behavioral paradigm and the delivery of a reward (sugary water) based on the success or failure of a visual task. Experiments were controlled by EventIDE software (Okazolab, Netherlands). Primates were under mild fluid restriction (approximately 30 mL/kg/day) and could drink ad libitum while working.

### Behavioral paradigm for SEF data

Monkey S was trained to perform an active oculomotor task consisting of successive and randomized pro-saccades and antisaccades. After a baseline (100 to 120 seconds, random), the monkey had to perform a pro-saccade (look at the presented target) if the presented cue was a vertical rectangle or an anti-saccade (look on the opposite side of the presented target) if the above-mentioned rectangle was horizontal. After a successful trial, the monkey received a sugary water reward (few drops) and an intertrial of 3 to 4 seconds (time between the end of a trial and the beginning of the next trial) was applied. If the trial was wrong, no sugary water was delivered and the intertrial started. The monkey worked for approximately one hour.

### Behavioral paradigm for V1 data

Monkey L was trained to perform passive fixation tasks. All sessions (except two) began with a 100s-baseline during which no stimuli were presented. The animal started the trial by fixing a central green square subtending 0.2 degrees of visual angle (DVA) for a random duration between 500 and 1000 ms within a tolerance window of 1.5 DVA. A visual stimulus of the peripheral location was then presented for 1s. We used different types of stimuli among sessions that were all localized in the left visual field. For most sessions, a single stimulus was presented. First, we used hemi-concentric bands of 2 DVA width centered on the central fixation point, filled with sinusoidal gratings with a fixed temporal frequency (one cycle per degree). The eccentricity was locked to 6 DVA. Second, we used polar angle stimuli that were 15 DVA of angular width extending from 1.5 to 15 DVA filled with sinusoidal gratings with a fixed temporal frequency (one cycle per degree). Third, we used gratings with varying spatial frequencies (3, 6, 9, 15 cycles per degree) in a rectangle area that covered the bottom-left quarter of visual field (DVA: -8 to 0 in the X axis, -7 to 0 in the Y axis). Fourth, we used a local checkerboard of 4*4 DVA center at R=6 DVA and theta = -¾π radians. The animal was rewarded by a small drop of liquid (sugary water) at the end of each correct fixation trial. We imposed an intertrial interval of 9 s to ensure that the CBV value came back to the initial value. We also used control trials with the same temporal organization but without any peripheral visual stimulus. All conditions were randomly interleaved.

### Implant and probe for functional ultrasound imaging For Awake Cooperative Monkeys

The head of the monkey was fixed using a surgically implanted titanium head post (Crist Instrument, MD, USA). After behavioral training of the animals, a recording chamber (CILUX chamber, Crist Instrument, MD, USA) was implanted and a craniotomy (diameter 19 mm) was performed (for monkey S: mediolateral: +0mm, anteroposterior: +26mm; for monkey L: mediolateral: +7 mm, anteroposterior: -10 mm, dorso-ventral: +38 mm). A custom ultrasonic probe (128 elements, 15 MHz, 100 × 100 μm² of spatial resolution) with sterile ultrasonic gel was used in the chamber. The acquired images had a pixel size of 100 × 100 μm and a slice thickness of 400 μm. We could image 12 mm along the cortical surface and up to 15 mm in depth.

### Functional ultrasound (fUS) recordings

Changes in CBV were measured using a real time functional ultrasound scanner prototype (Iconeus and Inserm U1273, Paris, France) with a custom 15-MHz linear probe. Data were acquired by emitting continuous groups of 11 planar ultrasonic waves tilted at angles varying from -10° to 10°. Ultrasonic echoes were summed to create a single compound image acquired every 2 ms. Final Doppler images were sampled at 2.5Hz by averaging 200 compound ultrasonic images after spatiotemporal filtering based on the singular value decomposition of the ultrasonic images.

### Eye movements and pupil recordingss

Eye movements and pupil diameter were recorded during the tasks using a video eye tracker (Eyelink 1k, SR-Research) connected to an analog-to-digital converter (Plexon Inc, TX, USA). All data were collected using Plexon software and analyzed using MATLAB (The MathWorks Inc.,Massachusetts, USA). Saccades were detected when the eye’s horizontal velocity went over 30°/.s.

#### Online microelectrode position monitoring

We slowly (5 µm.s-1) inserted a microelectrode through the cortex via the adapter tube while performing live fUS imaging of the region of interest. The microelectrode is not visible in ultrasonic images and its location in the imaged plane is obtained by estimating local motion thanks to the Singular Value Decomposition (SVD) of the ultrasonic raw data. Because the electrode moved slowly, we adjusted the SVD clutter filter from λ = 30 to λ = 2 to track the electrode movement accurately rather than blood flow. We determined the optimum value for the SVD clutter filter in NPH using the adaptive spatiotemporal SVD as described by Baranger et al.^53^ and determined λ = 30 as the optimum SVD clutter filter for the discrimination between tissue motion and blood flow in NHP. However, as the electrode is moving slower than red blood cells in vessels, we changed the clutter filter to isolate the insertion motion signal from the electrode and experimentally determined λ = 2 as the optimal value for the SVD clutter filter for online electrode monitoring.

### Extracellular electrophysiological recordings

Extracellular neuronal activity and local field potential (LFP) were recorded in vivo simultaneously with fUS recording, using tungsten microelectrodes of impedance ranging from 8 to 10 MΩ for monkey S (ref UEWLGASEFN1E, FHC Inc, ME, USA) or glass-coated tungsten electrodes of impedance ranging from 0.5 to 2.5 MΩ for monkey L (reference 366-120615-00, Alpha Omega, USA). The microelectrode was inserted into a 60 to 70mm stainless steel tubing and the tip of the microelectrode was connected to an amplifier with a male gold-plated pin connector (Ref. #520200, A-M Systems, WA, USA). The extracellular signal was amplified and collected using Plexon software. Offline spike analysis was performed manually using Plexon Offline Sorter (Plexon Inc, TX, USA). The microelectrode motion into cortical layers was performed using a micro-descender.

### Setup preparation

The bottom of the recording chamber was filled with sterile ultrasonic gel. The ultrasonic probe and microelectrode with steel guide tube were inserted into the designed co-recording device. Hereafter, the device was inserted into the recording chamber, until contact with the ultrasonic gel. Appropriate position of the device was verified by fUS live view acquisition.

### fUS data processing

Doppler data were analyzed using a generalized linear model approach implemented in Matlab. The stimulation pattern, consisting of a Dirac comb with all cues presentations, in the design matrix was convoluted with the fUS-determined hemodynamic response function and a Z-score and p-value map were obtained. The activation maps show the Z-score of all pixels in the images with a p-value < 0.001 (before Benjamini-Hochberg procedure to decrease false discovery rate). We chose the region of interest (ROI) within the supplementary eye field based on the Z-score map and Paxinos atlas for macaque brains and the signal was averaged to obtain a single temporal signal. The spatially averaged signal was then expressed as the relative increase in CBV (in percent) by subtracting the baseline CBV (calculated during the baseline at the beginning of an acquisition) followed by division of the difference by the baseline CBV. Same analysis was run for V1.

### Processing of electrophysiological signals

Electrophysiological data were spike-sorted both online and offline, using Plexon software (Plexon Offline Sorter v4.5.1, Plexon, Dallas, TX, USA) during and after data acquisition. After the spike sorting, spikes are realigned on different time events (time of the cue presentation, time of the saccade for monkey S. and time of the reward) to determine the nature of the recorded neuron and the spike density function (SDF). The spike-density function was produced by convolving the spike train from each trial with a function resembling a postsynaptic potential specified by τg, the time constant for the growth phase, and τd, the time constant for the decay phase as *R*(*t*) = (1 − exp(−*t*/τd)*exp(−*t*/τd). Based on physiological data from excitatory synapses τg was set to 1 ms and τd to 20 ms / 50ms for Monkey S and Monkey L, respectively (Sayer et al. 1990). The magnitude of the visual response was determined for each cell as the maximum value of the spike-density function during the time interval between the onset and the end of visual response. For the statistical parametric mapping, we needed an input signal linked to the neural activity. Here, we simply convolved spikes trains with the haemodynamic response function determined for each animal for their respective studied brain region (SEF for Monkey S. and V1 for Monkey L.) sampled at 1kHz and downsampled this signal to fUS imaging rate (2.5 Hz) before applying the generalized linear model for the statistical parametric mapping algorithm.

### Cross-correlation between signals

In order to calculate the cross-correlation between fUS signal and spikes density function, we down-sampled the SDF to fUS signal frequency (2.5Hz). Signals were then cut in non-overlapping 10 seconds windows and cross-correlations were calculated using MATLAB *xcorr* function with *‘normalized’* option (in order to normalize signals in such a way that auto-correlation at null lag is equal to 1). Same protocol was used to calculate fUS-fUS cross-correlation. Afterwards, in order to have the cross-correlation between our signals through time, we took the cross-correlation at t = + 2 sec for fUS-SDF and t = 0 sec for fUS-fUS cross-correlation. As we wished to compare the cross-correlation during the baseline and the cross-correlation during the task, we used *paired T-test* after verifications of data normality using the Shapiro-Wilk test.

---

**Supplementary Figure 1.**
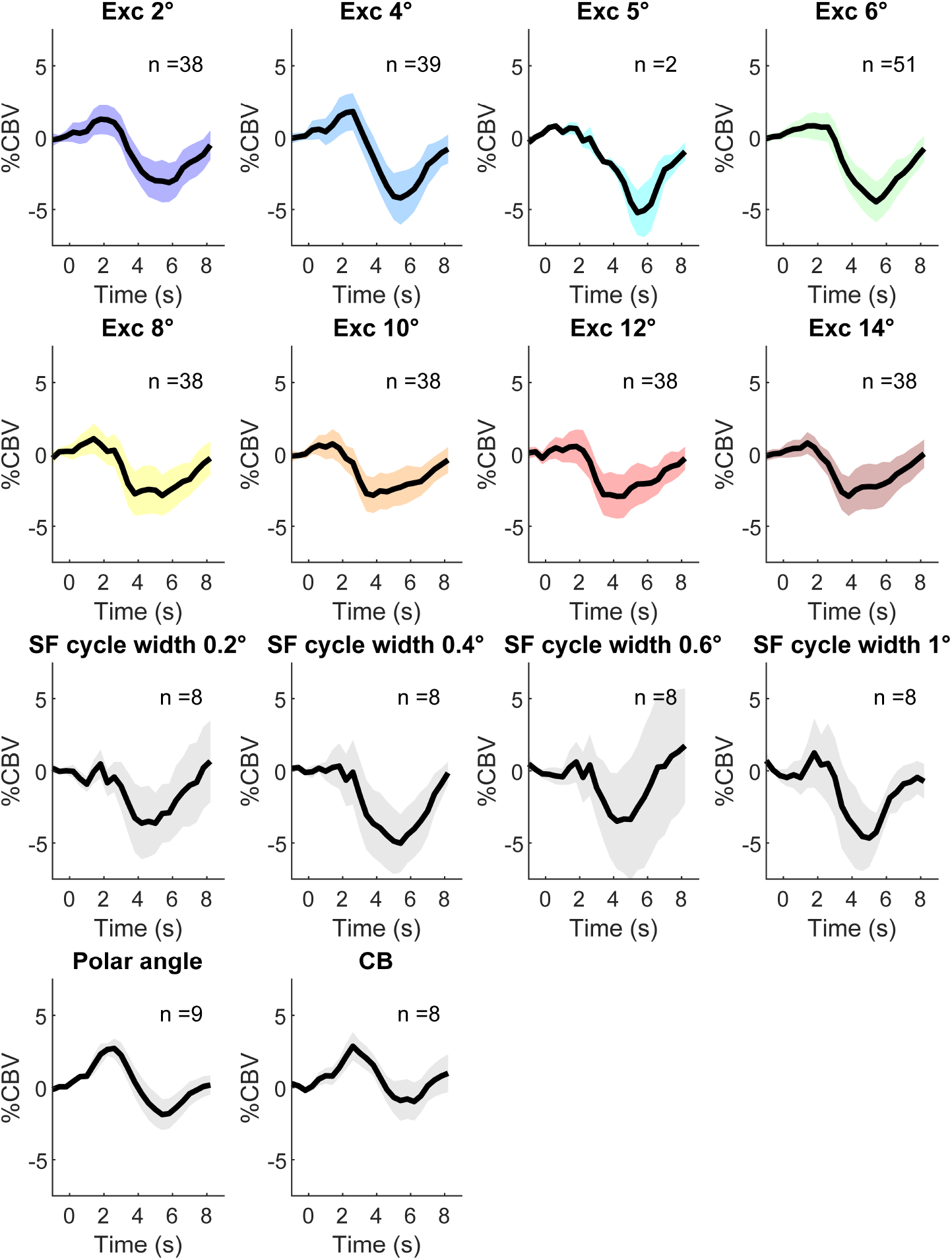
Mean CBV variations among sessions for each stimulus used. Shaded areas represent standard deviation. The number of sessions per stimulus is indicated (n=79 in total). ‘Exc’ refers to hemi-concentric bands of 2 DVA width with varying eccentricities. ‘SF’ refers to gratings with varying spatial frequencies in a rectangle area that covers the bottom-left quarter of visual field (DVA : -8 to 0 in the X axis, -7 to 0 in the Y axis). ‘Polar angle’ refers to the polar angle stimulus with 15 DVA of angular width extending from 1.5 to 15 DVA. ‘CB’ refers to local checkerboard.

**Supplementary Figure 2.**
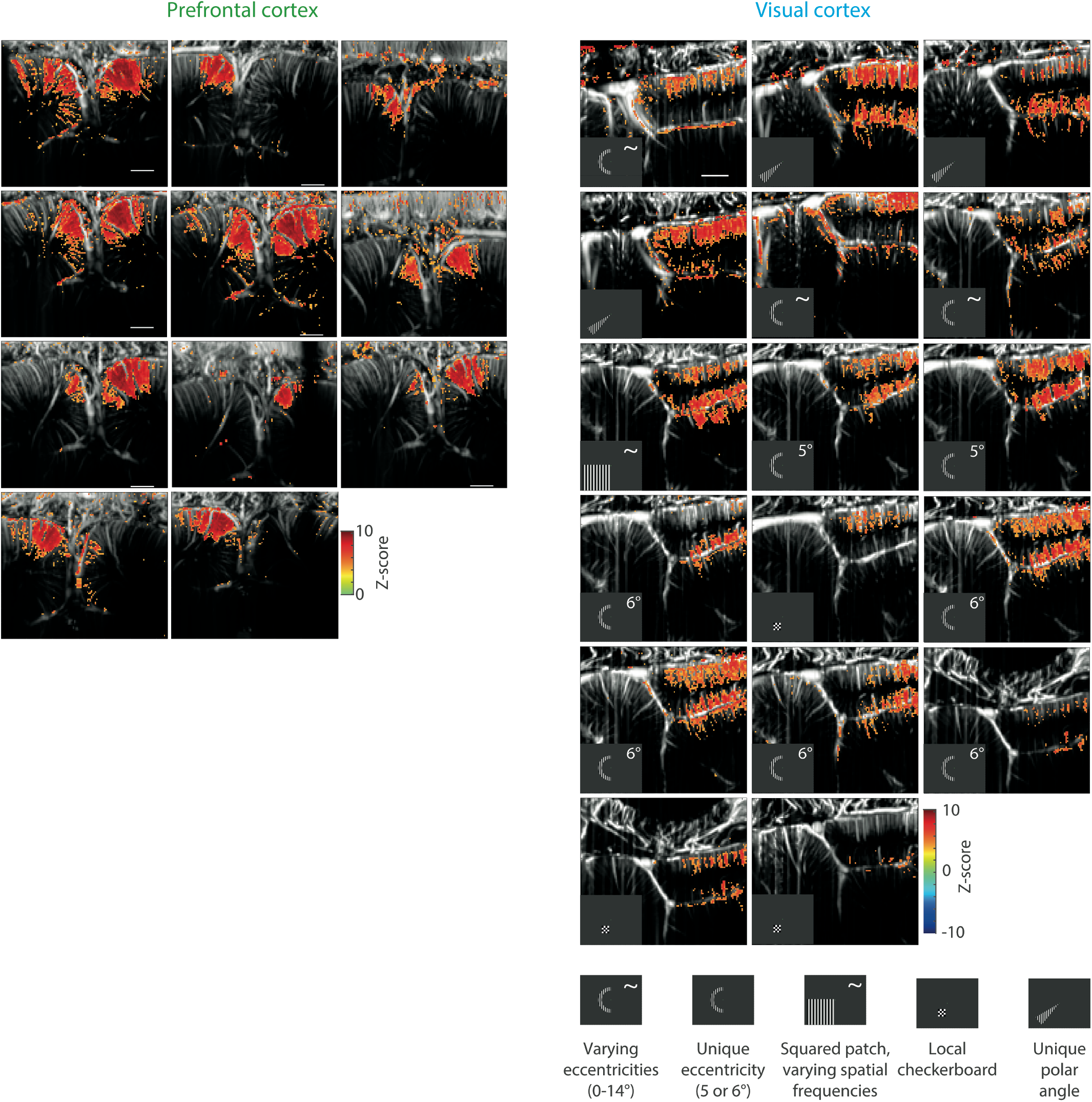
All GLM stimuli-induced maps for SEF and V1. Stimuli used for each session for V1 are indicated.

**Supplementary Figure 3.**
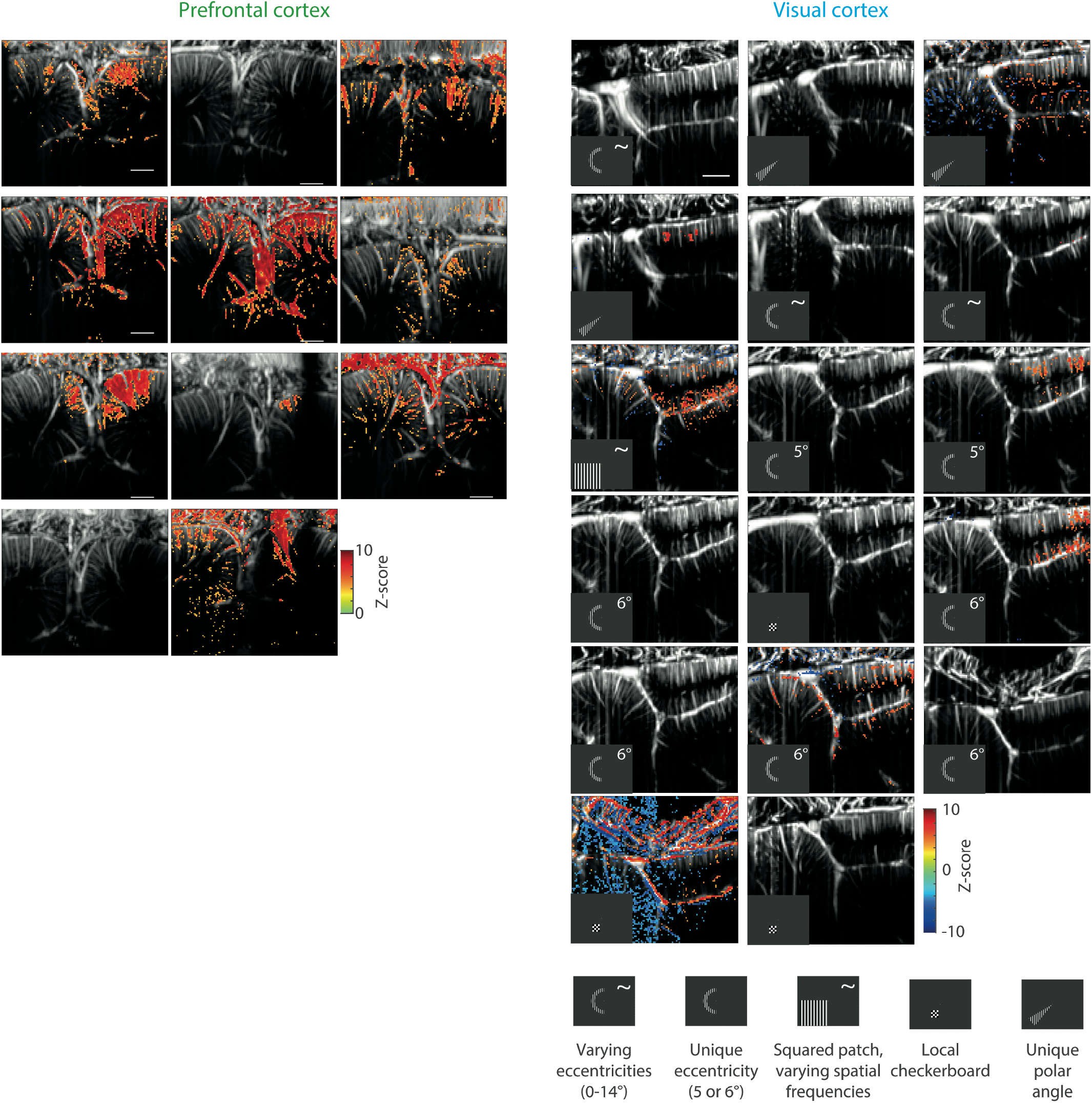
All GLM spikes-induced maps for SEF and V1. Stimuli used for each session for V1 are indicated.

